# Kinetochore phosphatases suppress autonomous kinase activity to control the spindle assembly checkpoint

**DOI:** 10.1101/856773

**Authors:** Marilia H Cordeiro, Richard J Smith, Adrian T Saurin

## Abstract

Local phosphatase regulation is critical for determining when phosphorylation signals are activated or deactivated. A typical example is the spindle assembly checkpoint (SAC) during mitosis, which regulates kinetochore PP1 and PP2A-B56 activities to switch-off signalling events at the correct time. In this case, kinetochore phosphatase activation dephosphorylates MELT motifs on KNL1 to remove SAC proteins, including the BUB complex. We show here that, surprisingly, neither PP1 or PP2A are required to dephosphorylate the MELT motifs. Instead, they remove polo-like kinase 1 (PLK1) from the BUB complex, which can otherwise maintain MELT phosphorylation in an autocatalytic manner. This is their principle role in the SAC, because both phosphatases become redundant if PLK1 is inhibited or BUB-PLK1 interaction is prevented. Therefore, phosphatase regulation is critical for the SAC, but primarily to restrain and extinguish autonomous kinase activity. We propose that these circuits have evolved to generate a semi-autonomous SAC signal that can be synchronously silenced following kinetochore-microtubule tension.

## Introduction

The spindle assembly checkpoint (SAC) prevents mitotic exit until chromosomes have attached to microtubules via the kinetochore ^1,2^. MPS1 kinase initiates SAC signalling by localising to unattached kinetochores and phosphorylating the SAC scaffold KNL1 on repeat motifs known as ‘MELT repeats’ (for the amino acid consensus Met-Glu-Leu-Thr) ^3–5^. Once phosphorylated, these MELT motifs recruit the heterotetrameric BUB1-BUB3-BUB3-BUBR1 complex (hereafter BUB complex) to kinetochores ^6–9^, which, directly or indirectly, recruits all other proteins needed to activate the SAC and block mitotic exit ^1,2^. Once kinetochores attach to microtubules, the local SAC signal must be rapidly extinguished by at least three different mechanisms: 1) localised MPS1 activity is inhibited ^10–12^, 2) key phosphorylation sites, such as the MELT repeats, are dephosphorylated by KNL1-localised phosphatases ^13–17^, and 3) dynein motors physically transport SAC components away from kinetochores down microtubules ^18^.

One key unexplained aspect of SAC signalling concerns the role of polo-like kinase 1 (PLK1) ^19^. PLK1 interacts via its polo-box domains (PBDs) to phospho-epitopes on various different kinetochore complexes, including to two CDK1 phosphorylation sites on the BUB complex (BUB1-pT609 and BUBR1-pT620) ^20–22^. PLK1 has similar substrates preferences to MPS1 ^23,24^ and it shares at least two key substrates that are critical for SAC signalling: the KNL1-MELT motifs and MPS1 itself, including key sites in the MPS1 activation loop ^25–27^. PLK1 can therefore enhance MPS1 kinase activity and also directly phosphorylate the MELT motifs to support SAC signalling, perhaps from its localised binding site on BUB1 ^27^. It is unclear why PLK1 is needed to cooperate with MPS1 in SAC signalling and, importantly, what inhibits PLK1 signalling to allow MELT dephosphorylation and SAC silencing upon microtubule attachment.

Our results demonstrate that two KNL1-localised phosphatases – PP1-KNL1 and PP2A-B56 –antagonise PLK1 recruitment to the BUB complex. This is crucial, because otherwise the BUB-PLK1 module primes further BUB-PLK1 recruitment to sustain the SAC in an autocatalytic manner. In fact, this is the primary role of both phosphatases in the SAC, because when PLK1 and MPS1 are inhibited together, MELTs dephosphorylation and SAC silencing can occur normally even when PP1 and PP2A are strongly inhibited. Therefore, this study demonstrates that the main role of KNL1-localised phosphatases is to suppress a semi-autonomous SAC signal that is driven by MPS1, but sustained by PLK1 and positive feedback.

## Results

### PP1-KNL1 and PP2A-B56 antagonise PLK1 recruitment to the BUB complex

Inhibition of PP1-KNL1 or knockdown of PP2A-B56 both enhance PLK1 recruitment to kinetochores ^28,29^. To test whether this was due to localised phosphatase inhibition at the BUB complex, we inhibited the recruitment of PP2A-B56 to BUBR1 (BUBR1^Δpp2A^) and compared this to a PP1-KNL1 mutant (KNL1^ΔPP1^), as used previously ^16,29^ (note, that in these and all subsequent experiments, siRNA-mediated gene knockdown was used in combination with doxycycline-inducible replacement of the wild type or mutant gene from an FRT locus: see methods). This demonstrated that removal of either PP1 from KNL1 or PP2A-B56 from BUBR1 increased PLK1 levels at unattached kinetochores (**Figure 1A**). We suspected that the increased PLK1 was due to enhanced binding to the BUB complex, because depletion of BUB1 and/or BUBR1 also removes the BUBR1:PP2A-B56 complex from kinetochores, but this did not enhance PLK1 levels (**Figures S1A-C**). In fact, kinetochore PLK1 was considerably reduced when the whole BUB complex was removed (see siBUB1 + siBUBR1 in **Figures S1A-C**). PLK1 binds via its PBDs to CDK1 phosphorylation sites on BUB1 (pT609 ^21^) and BUBR1 (pT620 ^20,22^), therefore we raised antibodies to these sites and validated their specificity in cells (**Figure S1D**). Immunofluorescence analysis demonstrated that phosphorylation of both sites is enhanced at unattached kinetochores in KNL1^ΔPP1^ or BUBR1^ΔPP2A^ cells (**Figures 1B-C, S1E-F**). This is the reason that PLK1 kinetochore levels increase when PP2A is removed, because the elevated PLK1 levels can be attenuated by BUBR1-T620A mutation (**Figure 1D**) and completely abolished by additional mutation of BUB1-T609A (**Figure 1E**). Therefore, these data demonstrate that PP1-KNL1 and BUBR1-bound PP2A-B56 antagonise PLK1 recruitment to the BUB complex by dephosphorylating key CDK1 phosphorylation sites on BUBR1 (pT620) and BUB1 (pT609) (**Figure 1F**).

**Figure 1.**
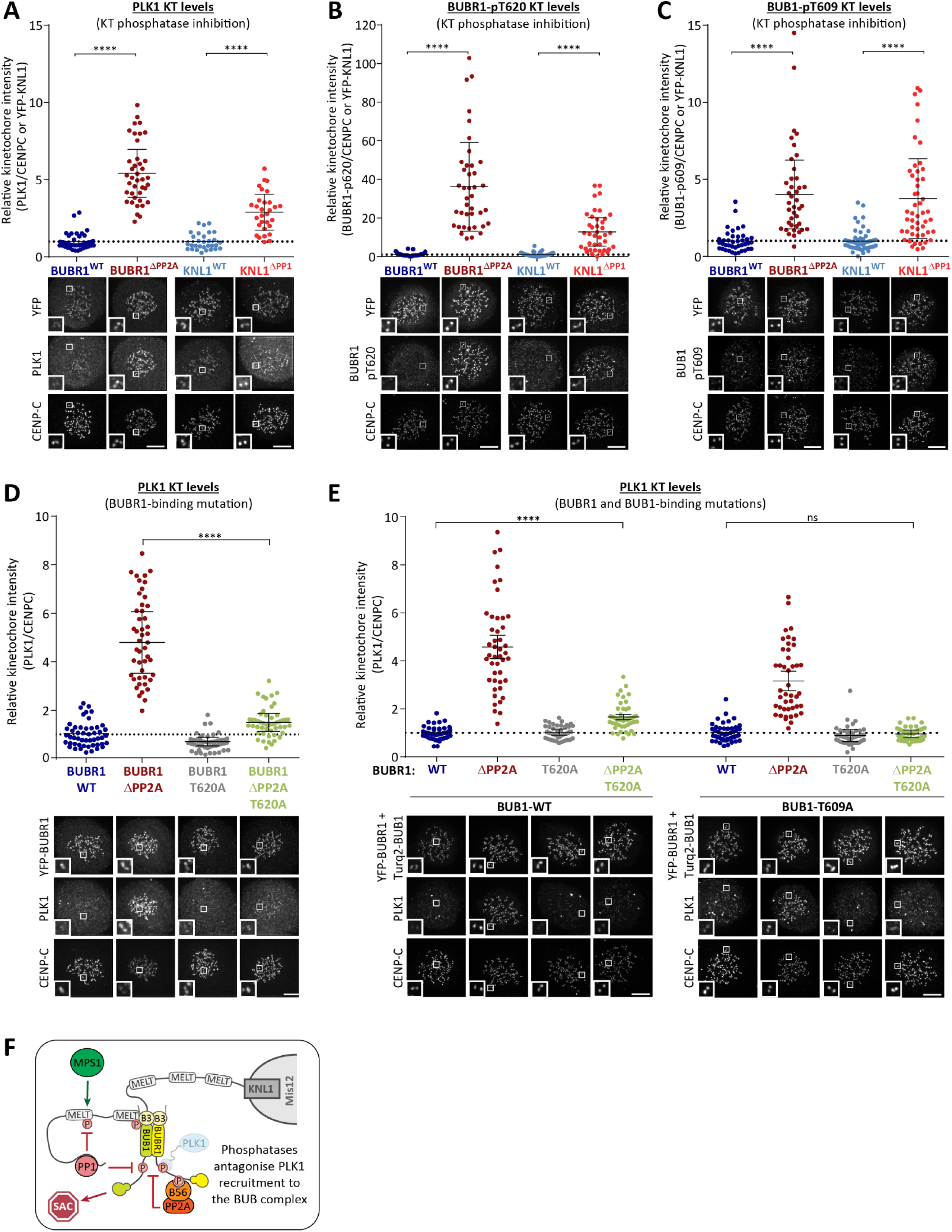
Kinetochore phosphatases PP1 and PP2A-B56 antagonise Plk1 recruitment to the BUB complex. **A-C)** Effect of phosphatase-binding mutants on levels of PLK1 (A), BUBR1-pT620 (B) and BUB1-pT609 (C) at unattached kinetochores in nocodazole-arrested cells. Mean kinetochore intensities from 30-50 cells, 3-5 experiments. **D-E)** Effect of mutating the PLK1-binding site on BUBR1 (pT620) (D) and BUB1 (pT609) (E) on PLK1 kinetochore levels in nocodazole-arrested BUBR1^WT/ΔPP2A^ cells. Mean kinetochore intensities from 40-50 cells per condition, 4-5 experiments.F) Model depicting how PP1 and PP2A-B56 dephosphorylate the BUB complex to inhibit PLK1 recruitment. For all kinetochore intensity graphs, each dot represents a cell, and the error bars display the variation between the experimental repeats (displayed as ± SD of the experimental means). Example immunofluorescence images were chosen that most closely resemble the mean values in the quantifications. The insets show magnifications of the outlined regions. Scale bars = 5μm. ****p<0.0001, ns = non-significant.

### PP1-KNL1 and PP2A-B56 antagonise PLK1 to allow SAC silencing

PLK1 is able to support SAC signalling by phosphorylating the MELT motifs directly ^25–27^. Therefore, we hypothesised that the increased BUB-PLK1 levels in KNL1^ΔPP1^ and BUBR1^ΔPP1^ cells could help to sustain MELT phosphorylation and the SAC in the absence of MPS1 activity. To address this, we examined the effect of PLK1 inhibition with BI-2536 under these conditions. **Figures 2A-B and S2A** show that MELT dephosphorylation and mitotic exit are both attenuated following MPS1 inhibition in nocodazole-arrested KNL1^ΔPP1^ or BUBR1^ΔPP1^ cells, as demonstrated previously ^16,17^.

**Figure 2:**
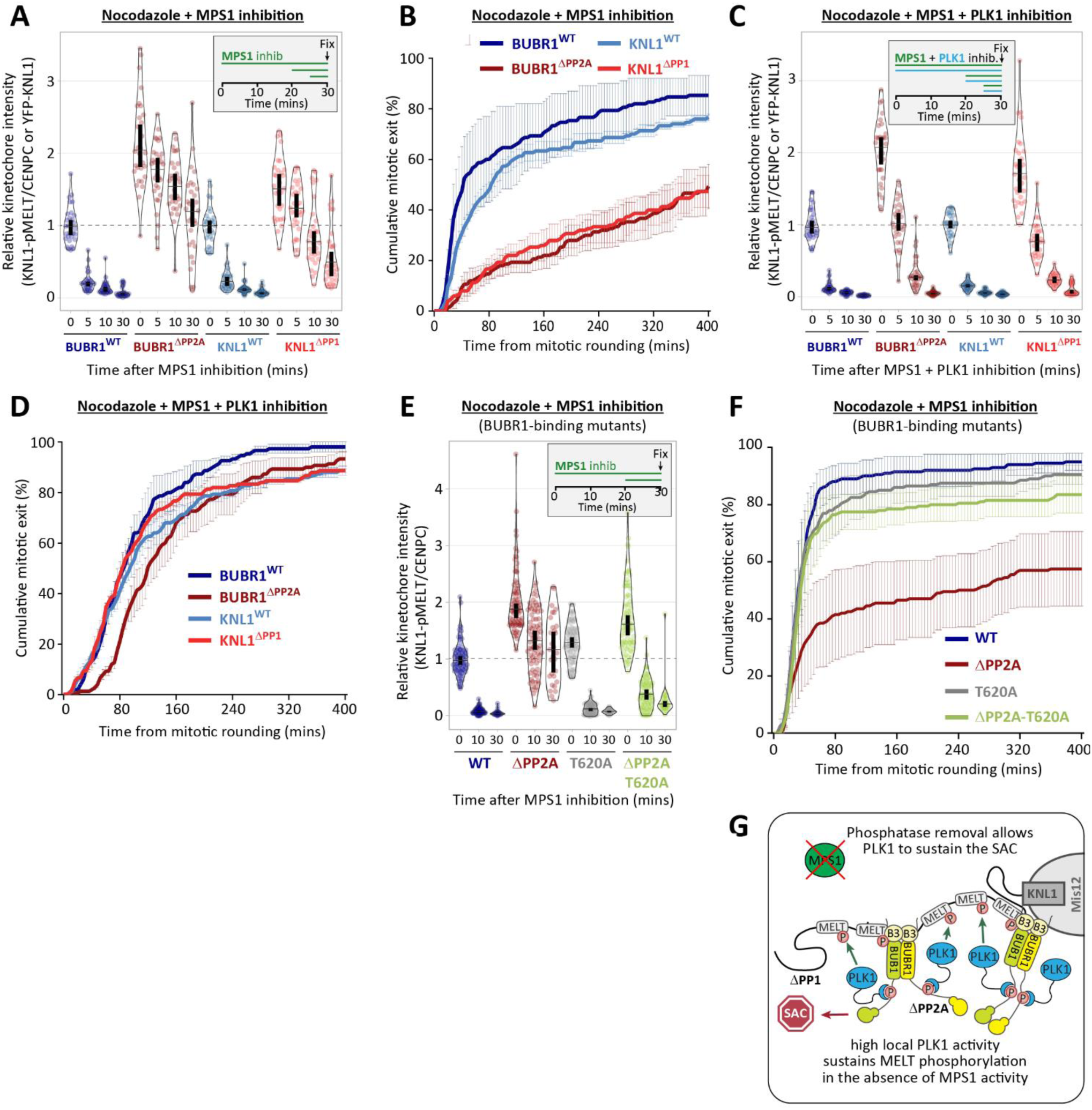
Kinetochore phosphatases PP1 and PP2A-B56 remove Plk1 from the BUB complex to silence the SAC. **A and C)** Effects of phosphatase-binding mutants on KNL1-MELT dephosphorylation in nocodazole-arrested cells treated with MPS1 inhibitor AZ-3146 (2.5 μM) for the indicated times, either alone (A) or in combination with the PLK1 inhibitor BI-2536 (100 nM) (C). MG132 was included in all treatments to prevent mitotic exit after addition of the MPS1 inhibitor. Graphs in A and C display kinetochore intensities from 30 cells per condition, 3 experiments. **B and D**) Effects of phosphatase-binding mutants on duration of mitotic arrest in nocodazole-arrested cells treated with MPS1 inhibitor AZ-3146 (2.5uM) alone (B) or in combination with PLK1 inhibitor BI-2536 (100nM) (D). Graphs in B and D show mean (± SD) of 150 cells per condition from 3 experiments. **E**) Effect of mutating the PLK1-binding site on BUBR1 (pT620) on MELT dephosphorylation (E) and mitotic exit (F) in nocodazole-arrested BUBR1^WT/ΔPP2A^ cells, treated as in A and B. Graph in E displays kinetochore intensities of 30-80 cells per condition from 3-7 experimental repeats. Graph in F shows the means (± SD) of 200 cells from 4 experiments. **G**) Model of how high PLK1 sustains MELT phosphorylation in the absence of MPS1 activity when kinetochore phosphatases are absent. In all kinetochore intensity graphs, each dot represents the mean kinetochore intensity of a cell, and the violin plots shows the distribution of intensities between cells. The thick vertical lines represent 95% confidence interval (CI) around the median, which can be used for statistical comparison of multiple timepoints/treatments by eye (see methods). Timelines indicate treatment regime prior to fixation.

Importantly, however, these effects are rescued if PLK1 and MPS1 are inhibited together (**Figures 2C-D and S2B**) (note, the thick vertical bars in these violin plots display 95% confidence intervals, which can be used for statistical comparison of multiple timepoints/treatment by eye: see methods). Therefore, when kinetochore phosphatase recruitment is inhibited, PLK1 becomes capable of supporting the SAC independently. This is due to enhanced PLK1 levels at the BUB complex, because MELT dephosphorylation and SAC silencing are also rescued if MPS1 is inhibited when the PLK1 binding motif on BUBR1 is mutated (BUBR1^ΔPP2A-T620A^, **Figure 2E-F**). Collectively these data demonstrate that excessive PLK1 levels at the BUB complex can sustain the SAC when KNL1-localised phosphatases are removed (**Figure 2G**). Therefore, these phosphatases promote SAC silencing by antagonising PLK1 recruitment to BUB1 and BUBR1. This raises the important question of whether PLK1 removal from the BUB complex is the *only* critical role for these phosphatases in the SAC, or whether they are additionally needed to dephosphorylate the MELT repeats, as previously assumed ^13–17^.

There is an approximate 5-10 min delay in MELT dephosphorylation between wild type and KNL1^ΔPP1^/BUBR1^ΔPP2A^ cells when MPS1 and PLK1 are inhibited together (blue vs red symbols in **Figure 2C**). This could indicate a role for PP1/PP2A in MELT dephosphorylation, or alternatively, it may reflect the time it takes for BI-2536 to penetrate cells and inhibit PLK1, since both inhibitors were added together at timepoint zero in this assay. Therefore, we next sought to dissect if localised PP1 or PP2A contribute directly to MELT dephosphorylation.

### PP1 and PP2A are not required for KNL1-MELT dephosphorylation

PLK1 inhibition for 30 minutes was sufficient to reduce phospho-MELT in ^ΔPP1^ and BUBR1^ΔPP1^ cells to levels comparable with wild type cells (**Figures 3A and S3A**). Therefore, we next examined MELT dephosphorylation rates when MPS1 was inhibited immediately after this 30-minute inhibition of PLK1. **Figures 3B and S3B** show that the MELT motifs are dephosphorylated with very similar kinetics in this assay, irrespective of whether PP1 or PP2A-B56 kinetochore recruitment is inhibited. This was particularly surprising, because it implies that neither PP1 or PP2A are essential for dephosphorylating the MELT repeats. This could reflect redundancy between the two phosphatases, therefore we attempted to remove both phosphatases from kinetochores by combining PP2A-B56 and PP1 knockdown (**Figure 3C**) or performing PP2A-B56 knockdown in KNL1^ΔPP1^ cells (**Figure 3D**). However, in both of these situations, MELT dephosphorylation was indistinguishable from wild type cells if MPS1 and PLK1 were inhibited together (**Figures 3C-D and S3C-F**). Finally, we treated cells with a high dose of Calyculin A (50 nM) - a very potent inhibitor of all PP1 and PP2A phosphatases (IC50 values, 1-2nM ^30^ – which can completely prevent MELT dephosphorylation and BUB complex removal following MPS1 inhibition alone (**Figures 3E and S3G-I**). However, even in this situation, the MELT motifs are still dephosphorylated and the BUB complex is removed with fast kinetics when MPS1 and PLK1 are inhibited together (**Figures 3E and S3G-I**).

Therefore, these data demonstrate that neither PP1 or PP2A are required to dephosphorylate the MELT motifs. Instead they are needed to remove co-localised PLK1 from the BUB complex. This is important because BUB1-bound PLK1 can otherwise prime its own recruitment (via pMELT) and therefore maintain the SAC signal in a semi-autonomous manner. **(Figure 4A)**

**Figure 3:**
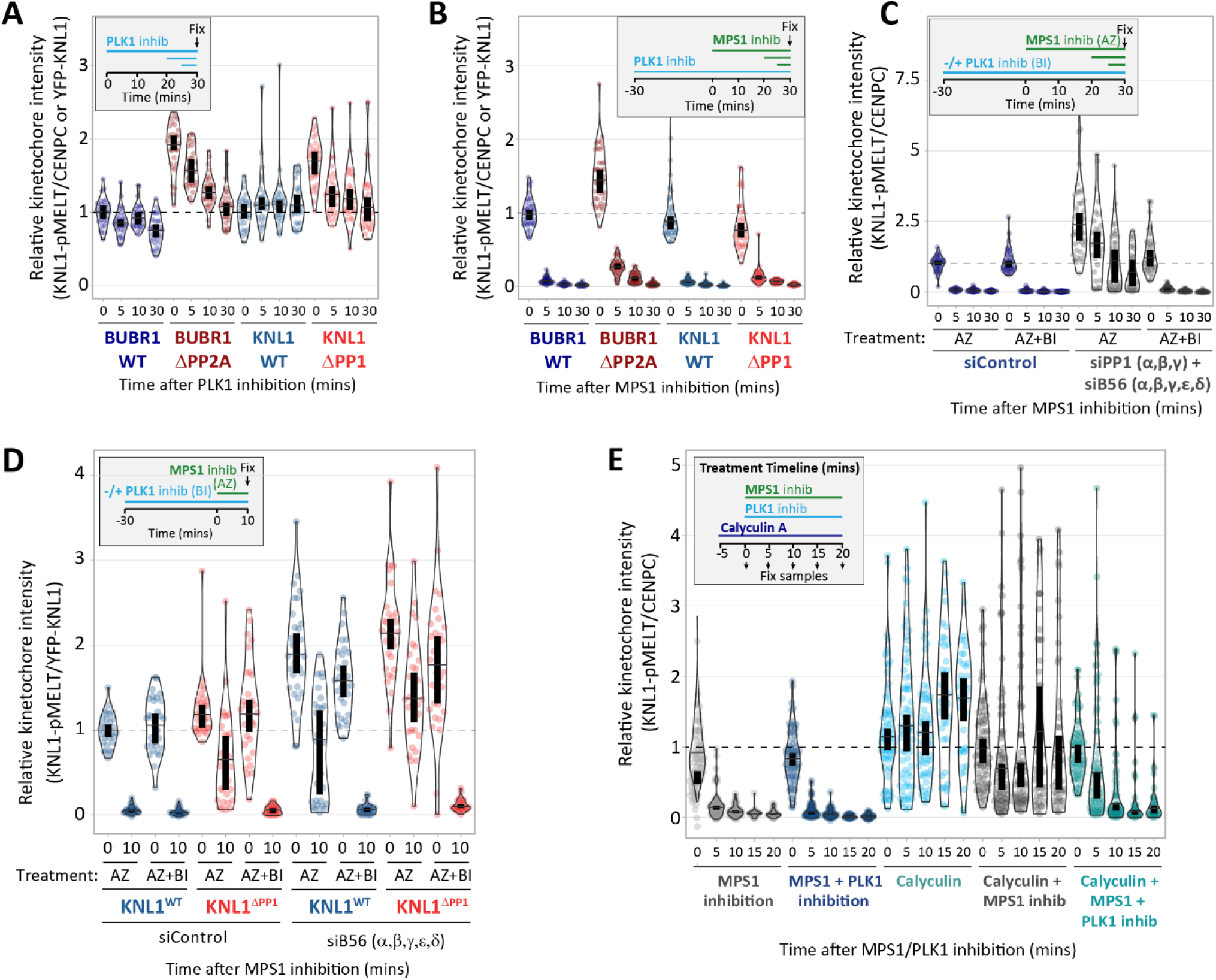
The main role of PP1 and PP2A-B56 in the SAC is to inhibit kinetochore-localised PLK1. **A-B)** Effects of phosphatase-binding mutants on KNL1-MELT dephosphorylation in nocodazole-arrested cells treated PLK1 inhibitor BI-2536 (100nM) alone (A) or in combination with MPS1 inhibitor AZ-3146 (2.5μM) (B), as indicated in the timelines. **C-D**) Knl1-MELT phosphorylation levels following combined siRNA-mediated knockdown of all PP1 and B56 isoforms (C) or all B56 isoforms in PP1^WT/ΔPP1^ cells (D). The quantifications are from nocodazole-arrested cells, treated with MPS1 inhibitor AZ-3146 (2.5μM) alone or in combination with PLK1 inhibitor BI-2536 (100 nM), as indicated. **E**) Knl1-MELT dephosphorylation in nocodazole-arrested cells treated with kinase inhibitors in the presence or absence of the PP1/PP2A phosphatase inhibitor calyculin A (50 nM), as indicated. In all kinetochore intensity graphs, each dot represents the mean kinetochore intensity of a cell, and the violin plots shows the distribution of mean intensities between cells. The thick vertical lines represent 95% confidence interval (CI) around the median, which can be used for statistical comparison of multiple timepoint/treatments by eye (see methods). Graphs in A-D derived from 30-40 cells per condition, 3-4 experiments. Graph in E derived from 50-70 cells, 5-7 experiments. Timelines indicate treatment regime prior to fixation. MG132 was included in combination with MPS1 inhibitor, in every case, to prevent mitotic exit.

**Figure 4:**
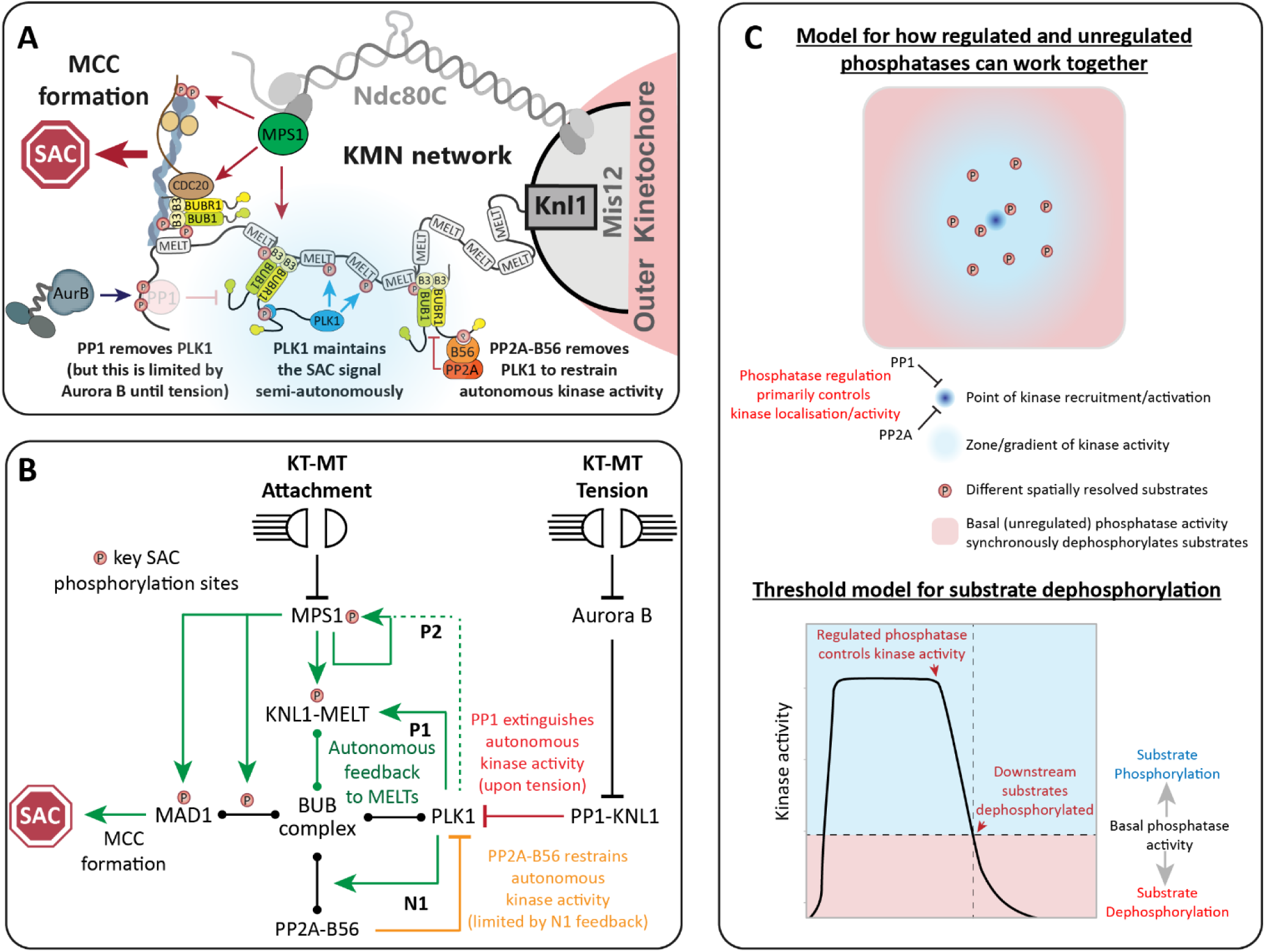
Kinetochore phosphatases restrain (PP2A) or extinguish (PP1) autonomous kinase activity to control the SAC. Schematic model to (**A**) summarise the results, and (**B**) illustrate the relevant feedback loops (see discussion for explanations). C)Model for how localised processes could use a combination of regulated and unregulated phosphatase activity to synchronously dephosphorylate spatially resolved substrates.

## Discussion

Serine/threonine phosphatases were once considered slaves to their kinase counterparts: kinases respond to regulatory inputs whereas phosphatases provide the basal level of activity needed for the kinases to act. A logical interpretation perhaps, considering that phosphatases display little specificity in vitro and are encoded by a relatively small number of genes that lack complex regulatory elements. However, this kinase-centric view of signalling changed when it became clear that phosphatases bind to substrates or adaptors in a regulated manner using specific short-linear motifs (SLiMs)^31^. In the case of PP1 and PP2A-B56, these SLiM interactions can be either inhibited (PP1) or enhanced (PP2A-B56) by phosphorylation inputs ^32,33^. We demonstrated recently that kinetochores use these phosphatases primarily because of their ability to be regulated in this way ^34^.

We build on this work here to show that PP1 and PP2A-B56 regulation is critical for the SAC, but primarily to determine when localised kinase activity is extinguished. The kinase, in this case, is PLK1 which is recruited via the BUB complex to the SAC scaffold KNL1. PLK1 is able to phosphorylate KNL1 on MELT motifs to recruit further BUB complexes, which, directly or indirectly, recruits all other proteins needed to generate the SAC signal ^1,2,25–27^. Therefore, this positive feedback loop helps to maintain the SAC in a semi-autonomous manner (pMELT→BUB:PLK1→pMELT, **Figure 4B, P1**). Multiple MELT repeats are active on each KNL1 molecule ^8,9,35^ and multiple copies of KNL1 are present on each kinetochore ^36^. Therefore, this creates over a thousand active MELT repeats per kinetochore, which the BUB-PLK1 module can use to rapidly amplifying SAC signalling downstream of MPS1. Furthermore, this may be reinforced by an additional positive feedback loop to MPS1 itself, since PLK1 can phosphorylate the activation loop of MPS1 to stimulate its kinase activity ^26,27,37^ (MPS1→BUB:PLK1→MPS1, **Figure 4B, P2**). This may help to explain the recent observation that this ‘autoactivation’ site remains phosphorylated when MPS1 is inhibited in BUBR1^ΔPP1^ cells ^38^. In this case, PP2A removal may increase phosphorylation indirectly via PLK1, instead of (or in addition to) preventing dephosphorylation directly, as the authors propose. In summary, positive feedback downstream of PLK1 explains why this kinase cooperates with MPS1 to activate the SAC: it allows the SAC signal to be establish in a rapid, semi-autonomous, manner. This also explains why phosphatase regulation is critical to prevent these positive feedback loops from locking the SAC signal into a constitutive ‘on’ state.

An important phosphatase in this regard is PP2A-B56, which is recruited to the BUB complex by PLK1 itself ^20,39–41^. This create a key negative feedback loop which allows PLK1 to restrain its own activity (PLK1→PP2A⊣PLK1, **Figure 4B, N1).** This loop is crucial to dampen the SAC signal, because when PP2A-B56 is removed, PLK1 recruitment is increased and the SAC can be sustained in a manner that is dependent on PLK1, but largely independent of upstream MPS1 activity (**Figure 2**). We hypothesise that this loop also restrains PP2A-B56 activity to prevent it from fully removing PLK1 to silencing the SAC on its own. In support of this hypothesis, PP2A-B56 can shut down the SAC if it is recruited to BUBR1 in a manner that is independent of phosphorylation ^34^. This homeostatic SAC regulation by phospho-dependent PP2A may be important to preserve the BUB complex at kinetochores when microtubules attach and MPS1 is removed/inhibited ^10–12^. The benefit of preserving the SAC platform in this situation is that SAC signalling can be rapidly re-established if microtubules subsequently detach. However, if the correct bioriented stated is achieved and the kinetochore comes under tension, then the PP1 arm is engaged ^42^, which crucially, is not restricted by negative feedback (**Figure 4B**). In fact, we speculate that this tension-dependent switch is reinforced by positive feedback instead, because removal of PLK1 and the BUB complex may reduce the inhibitory Aurora B signal emanating from centromeres ^19, 43^.

In summary, by interrogating the specific role of PP1 and PP2A-B56 at kinetochores, we arrive at the conclusion that phosphatase regulation is critical, but primarily to restrain and extinguish localised kinase activities. The phosphatases are the metaphorical slaves once again, but this time, they live up to their billing because they are directly controlled by their kinase masters (i.e. by SLiM phosphorylation ^39–42^). This is important because it is ultimately the kinases that sense the microtubule attachment state of each kinetochore, before relaying that information to the phosphatases to determine whether the SAC signal stays on or shuts off (**Figure 4B**). In fact, in this final act, it is the slaves that are ultimately freed to silence their kinase masters. It is interesting to note that other organisms have evolved different circuits to control the SAC, but even in these cases, phosphatase regulation appears to focus back to limit kinase activity. In flies, the KNL1-MELTs are not phospho-regulated and MPS1/PLK1 activities are not required to recruit the BUB complex to kinetochores ^44,45^. However, in this situation, PP1 binds directly to MPS1 to inhibit its activity and silence the downstream SAC signal ^46^. Therefore, the use of regulated phosphatases to silence local kinase activity may be a conserved feature of SAC signalling.

One final important point concerns the fact that the ultimate dephosphorylation of SAC substrates, such as the MELT motifs, does not appear to depend on either PP1 or PP2A (**Figure 3**). Instead, we speculate that it relies on a constitutive phosphatase(s) that has unregulated basal activity. One advantage of using a constitutive phosphatase in this situation, is that it can reverse different signals in a synchronous manner, irrespective of their positions within the kinetochore. This may be difficult to achieve if the executioner phosphatase is a regulated one that binds to a defined location within one protein. The final general model is depicted in **Figure 4C**. The basic premise of this model is that some cellular processes, such as the SAC, use kinases to phosphorylate different spatially resolved substrates. When those substrates must be synchronously dephosphorylated in a regulated manner, then phosphatase regulation can extinguish kinase activity at its point of origin, before basal phosphatase activity dephosphorylates the substrates (when kinase activity falls below a certain threshold). It will be important to test this hypothesis in future and to determine whether other signalling pathways use a combination of regulated and unregulated phosphatase activity to shut down localised processes in a rapid synchronous manner. If phosphatase regulation has generally evolved to restrain or extinguish localised kinase activity, as we shown here for the SAC, then this regulation could be therapeutically targeted to specifically elevate kinase activity in a range of conditions. Recent work has demonstrated that inhibition of a specific regulated phosphatase complex is both achievable and therapeutically valuable ^47,48^.

## Acknowledgements

This work was funded by a Cancer Research UK Programme Foundation Award to ATS (C47320/A21229 and C10988/A22566). We thank staff at the Dundee Imaging Facility and the Genetic Core Services Unit. We also thank Stephen Taylor for providing the HeLa Flp-in cell line and Geert Kops for antibodies.

## AUTHOR CONTRIBUTIONS

ATS, RJS and MHC conceived the study, designed the experiments and interpreted the data. RJS and MHC performed the experiments. ATS supervised the study and wrote the manuscript with input from all authors.

## DECLARATION OF INTERESTS

The authors declare no competing interests.

## MATERIAL AND METHODS

### Cell culture

All cell lines were derived from HeLa Flp-in cells (a gift from S Taylor, University of Manchester, UK) ^49^ and authenticated by STR profiling (Eurofins). The cells were cultured in DMEM supplemented with 9% FBS and 50 μg/ml penicillin/streptomycin. During fluorescence time-lapse analysis, cells were cultured in DMEM (no phenol red) supplemented with 9% FBS and 50μg/ml penicillin/streptomycin. Cells were screened every 4-8 weeks to ensure they were mycoplasma free.

### Plasmids and cloning

pcDNA5-YFP-BUBR1^WT^ expressing an siRNA-resistant and N-terminally YFP-tagged wild-type BUBR1 and pcDNA5-YFP-BUBR1^ΔPP2A^ (also called BUBR1^ΔKARD^), lacking amino acids 663-680 were previously described ^16^. These vectors were used to generate pcDNA5-YFP-BUBR1^T620A^ and pcDNA5-YFP-BUBR1^ΔPP2A-T620A^ by site directed mutagenesis. pcDNA5-YFP-BUB1^WT^ expressing an siRNA-resistant and N-terminally YFP-tagged wild-type BUB1 was described previously ^50^ and used to generate pcDNA5-YFP-BUB1^T609A^ by site directed mutagenesis. pcDNA5-mTurquoise2(Turq2)-BUB1^WT^ and pcDNA5-Turq2-BUB1^T609A^ were created by restriction cloning using Acc65I and BstBI to replace the YFP originally present in pcDNA5-YFP-BUB1^WT^ and pcDNA5-YFP-BUB1^T609A^ (Turq2 subcloned from pcDNA4-mTurq2-BUBR1^WT 34^). pcDNA5-YFP-KNL1^WT^ expressing an siRNA-resistant and N-terminally YFP-tagged wild-type KNL1 and pcDNA5-YFP-KNL1^ΔPP1^ (with RVSF at amino acids 58-61 mutated to AAAA, also called KNL1^4A^) were previously described ^34^. All plasmids were fully sequenced to verify the transgene was correct.

### Gene expression

HeLa Flp-in cells were used to stably express doxycycline-inducible constructs after transfection with the relevant pcDNA5/FRT/TO vector and the Flp recombinase pOG44 (Thermo Fisher). Cells were then selected for stable integrants at the FRT locus using 200 μg/ml hygromycin B (Santa Cruz biotechnology) for at least 2 weeks. In experiment requiring two transgenes in Figure 1E, Turq2-BUB1^WT^ or Turq2-BUB1^T609A^ were transiently transfect into cells stably expressing doxycycline-inducible YFP-tagged BUBR1 mutants. Transfection with these Turq2-tagged constructs was done 32 hours prior to endogenous gene knock-down (described below) and at least 72 hours prior to fixation. Plasmids were transfected using Fugene HD (Promega) according to the manufacturer’s instructions.

### Gene knockdown

For all experiments in HeLa Flp-in cells, the endogenous mRNA was knocked down and replaced with an siRNA-resistant mutant. The siRNA’s used in this study were: BUBR1 (5’-AGAUCCUGGCUAACUGUUC −3’), siBUB1 (5’-GAAUGUAAGCGUUCACGAA −3’), B56α (PPP2R5A: 5’-UGAAUGAACUGGUUGAGUA −3’), B56β (PPP2R5B: 5’-GAACAAUGAGUAUAUCCUA −3’), B56γ (PPP2R5C: 5’-GGAAGAUGAACCAACGUUA −3’), B56δ (PPP2R5D: 5’-UGACUGAGCCGGUAAUUGU −3’), B56ɛ (PPP2R5E: 5’-GCACAGCUGGCAUAUUGUA −3’), PP1α (PP1CA: 5’-GUAGAAACCAUAGAUGCGG −3’), PP1β (PP1CB: 5’-ACAUCAGUAGGUCUCAUAA −3’), PP1γ (PP1CC: 5’-GCAUGAUUUGGAUCUUAUA −3’), GAPDH (control siRNA: 5’-GUCAACGGAUUUGGUCGUA−3’). All siRNA’s were synthesised with dTdT overhang by Sigma-Aldrich and used at 20nM final concentration (i.e. the pools for B56 or PP1 knockdown contain 20nM of each siRNA). Double stranded interference RNA was used to knockdown endogenous KNL1 (sense: 5’-GCAUGUAUCUCUUAAGGAAGAUGAA−3’; antisense: 5’-UUCAUCUUCCUUAAGAGAUACAUGCAU−3’) (Integrated DNA technologies) at a final concentration of 20 nM. All siRNAs/dsiRNAs were transfected using Lipofectamine^®^ RNAiMAX Transfection Reagent (Life Technologies) according to the manufacturer’s instructions.

After 16 h of knockdown, DNA synthesis was prevented by addition of thymidine (2mM, Sigma-Aldrich) for 24 h. Doxycycline (1μg/ml, Sigma-Aldrich) was used to induce expression of the BUBR1, BUB1 and KNL1 constructs during and following the thymidine block. Cells were then released from thymidine block into media supplemented with doxycycline and nocodazole (3.3 μM, Sigma-Aldrich) for 8.5 hours before processing for fixed analysis. In live-cell imaging experiments in Figure 2, MPS1 and/or PLK1 were inhibited by adding AZ-3146 (2.5 μM Sigma-Aldrich) and BI-2536 (100nM, Selleckbio) shortly prior to imaging (6-8h after thymidine release). For MPS1 and PLK1 inhibition in cells analysed by immunofluorescence, nocodazole and MG132 (10 μM, Sigma-Aldrich) were added first for 30 minutes (plus BI-2536 if pre-treatment is used), followed by a time-course of AZ-3146 and/or BI-2536 in media containing nocodazole and MG132.

A high dose of calyculin A (50nM, LC labs) was used to inhibit all PP1 and PP2A phosphatases (IC50 values, 1-2nM ^30^). In these experiments (Figures 3E and S3I), Hela FRT cells (empty FRT locus) were treated with nocodazole for 4 hours followed by a 30 min incubation with media containing nocodazole and MG132 (these two drugs were present in all the subsequent steps of these experiments) prior to treatment as indicated in the timelines.

To image nocodazole-arrested cells treated with siRNA targeting BUBR1, BUB1 or both (Figures S1A-C), HeLa FRT cells were released from thymidine block (40 h after siRNA treatment) for 7 hours before arresting at the G2/M boundary with RO-3306 treatment (10μM, Tocris) for 2 hours. Cells were then washed three times and incubated for 15 minutes with full growth media before addition of MG132 for 30 minutes to prevent mitotic exit. This is important so that cells enter mitosis in the presence of nocodazole and MG132, which allows the arrest to be maintained and Cyclin B levels to be preserved, even though the SAC is inhibited. Cells were then fixed and stained as described below.

### Immunofluorescence

Cells, plated on High Precision 1.5H 12-mm coverslips (Marienfeld), were fixed with 4% paraformaldehyde (PFA) in PBS for 10 min or pre-extracted (when using BUB1-pT609 or BUBR1-pT620 antibodies and in double mutant experiments) with 0.1% Triton X-100 in PEM (100 mM PIPES, pH 6.8, 1 mM MgCl_2_ and 5 mM EGTA) for 1 minute before addition of 4% PFA for 10 minutes. In experiments using calyculin A, coverslips were coated with poly-L-lysine to prevent cell loss (Sigma-Aldrich). After fixation, coverslips were washed with PBS and blocked with 3% BSA in PBS+ 0.5% Triton X-100 for 30 min, incubated with primary antibodies overnight at 4°C, washed with PBS and incubated with secondary antibodies plus DAPI (4,6-diamidino2-phenylindole, Thermo Fisher) for an additional 2-4 hours at room temperature in the dark. Coverslips were then washed with PBS and mounted on glass slides using ProLong antifade reagent (Molecular Probes). All images were acquired on a DeltaVision Core or Elite system equipped with a heated 37°C chamber, with a 100x/1.40 NA U Plan S Apochromat objective using softWoRx software (Applied precision). Images were acquired at 1×1 binning using a CoolSNAP HQ or HQ2 camera (Photometrics) and processed using softWorx software and ImageJ (National Institutes of Health). For experiments involving YFP-expressing cells, mitotic cells were selected based on good expression of YFP at the kinetochore (KNL1) or cytoplasm (BUBR1 cells). In cases were double mutants (YFP and Turquoise 2) were used (Figure 1E), cells were selected based on good YFP signal in the kinetochore and cytoplasm, since the chicken anti-GFP antibody used cannot discriminate between the two fluorescent proteins. All immunofluorescence images displayed are maximum intensity projections of deconvolved stacks and were chosen to most closely represent the mean quantified data.

### Time-lapse analyses

For fluorescence time-lapse imaging, cells were imaged in 24-well plates in DMEM (no phenol red) with a heated 37°C chamber in 5% CO_2_. Images were taken every 4 minutes with a 20x /0.4 NA air objective using a Zeiss Axio Observer 7 with a CMOS Orca flash 4.0 camera at 4×4 binning. For brightfield imaging, cells were imaged in a 24-well plate in DMEM in a heated chamber (37°C and 5% CO_2_) with a 10x/0.5 NA objective using a Hamamatsu ORCA-ER camera at 2×2 binning on a Zeiss Axiovert 200M, controlled by Micro-manager software (open source: https://micro-manager.org/) or with a 20x/0.4 NA air objective using a Zeiss Axio Observer 7 as detailed above. Mitotic exit was defined by cells flattening down in the presence of nocodazole and MPS1 inhibitor.

### Antibodies

All antibodies were diluted in 3% BSA in PBS. The following primary antibodies were used for immunofluorescence imaging (at the final concentration indicated): chicken anti-GFP (ab13970 from Abcam, 1:5000), guinea pig anti-Cenp-C (BT20278 from Caltag + Medsystems, 1:5000), rabbit anti-BUB1 (A300-373A from Bethyl, 1:1000), mouse anti-BUBR1 (A300-373A from Millipore, 1:1000), rabbit anti-PLK1 (IHC-00071 from Bethyl, 1:1000) and rabbit anti-BUBR1 (A300-386A from Bethyl, 1:1000). The rabbit anti-pMELT-KNL1 antibody is directed against Thr 943 and Thr 1155 of human KNL1 ^16^ (Gift from G.Kops, Hubrecht, NL). The rabbit anti-BUB1-p609 antibody was raised against phospho-Thr 609 of human BUB1 using the following peptide C-AQLAS[pT]PFHKLPVES (custom raised by Biomatik, 1:1000). The rabbit anti-BUBR1-p620 antibody was raised against phospho-Thr 620 of human BUBR1 using the following peptide C-AARFVS[pT]PFHE (custom raised by Moravian, 1:1000). Secondary antibodies used were highly-cross absorbed goat, anti-chicken, anti-rabbit, anti-mouse or anti-guinea pig coupled to Alexa Fluor 488, Alexa Fluor 568, or Alexa Fluor 647 (Thermo Fisher).

The following antibodies were used for western blotting (at the final concentration indicated): rabbit anti-GFP (custom polyclonal, a gift from G. Kops; 1:5000), rabbit anti-BUBR1 (A300-386A, Bethyl; 1:1000), rabbit anti-BUB1 (A300-373A from Bethyl, 1:1000), rabbit anti-BUB1-p609 (custom raised by Biomatik, 1:1000), rabbit anti-BUBR1-p620 (custom raised by Moravian, 1:1000) and mouse anti-α-Tubulin (clone B-5-1-2, T5168, Sigma-Aldrich, 1:10000). Secondary antibodies used were goat anti-mouse IgG HRP conjugate (Bio-Rad; 1:2000) and goat anti-rabbit IgG HRP conjugate (Bio-Rad; 1:5000).

### Immunoprecipitation and immunoblotting

Flp-in HeLa cells were treated with thymidine and doxycycline for 24 hours, followed by a treatment with nocodazole and doxycycline for 16 hours. Mitotic cells were isolated by mitotic shake off and incubated with media containing nocodazole and doxycycline with or without calyculin A (50nM, LC labs) for 20 min. Cells were lysed in lysis buffer (50 mM Tris pH 7.5, 150 mM NaCl, 0.5% TritonX-100, 1 mM Na_3_VO_4_, 5 mM ß-glycerophosphate, 25 mM NaF, 10 nM Calyculin A and complete protease inhibitor containing EDTA (Roche)) on ice for 20 minutes. The lysate was incubated with GFP-Trap magnetic beads (from ChromoTek) for 2 hr at 4°C on a rotating wheel in wash buffer (same as lysis Buffer, but without TritonX-100) at a 3:2 ratio of wash buffer:lysate. The beads were washed 3x with wash buffer and the sample was eluted according to the protocol from ChromoTek.

### Image quantification and statistical analysis

For quantification of kinetochore protein levels, images of similarly stained experiments were acquired with identical illumination settings and analysed using an ImageJ macro, as described previously ^51^. Two-tailed, non-parametric Mann-Whitney unpaired t-tests were performed to compare the means values between experimental groups in immunofluorescence quantifications from Figures 1 and S1 (using Prism 6 software). For Figure 2 onwards, when multiple timepoints and treatments are used to compare the difference in dephosphorylation kinetics, the graphs are plotted as violin plots using PlotsOfData (https://huygens.science.uva.nl/PlotsOfData/; ^52^). This allows the spread of data to be accurately visualised along with the 95% confidence intervals (thick vertical bars) calculated around the median (thin horizontal lines). to allow statistical comparison between all treatments and timepoints. When the vertical bar of one condition does not overlap with one in another condition the difference between the medians is statistically significant (p<0.05).

