## Supplementary Figures 1-3 for "Kinetochore phosphatases suppress autonomous kinase activity to control the spindle assembly checkpoint"

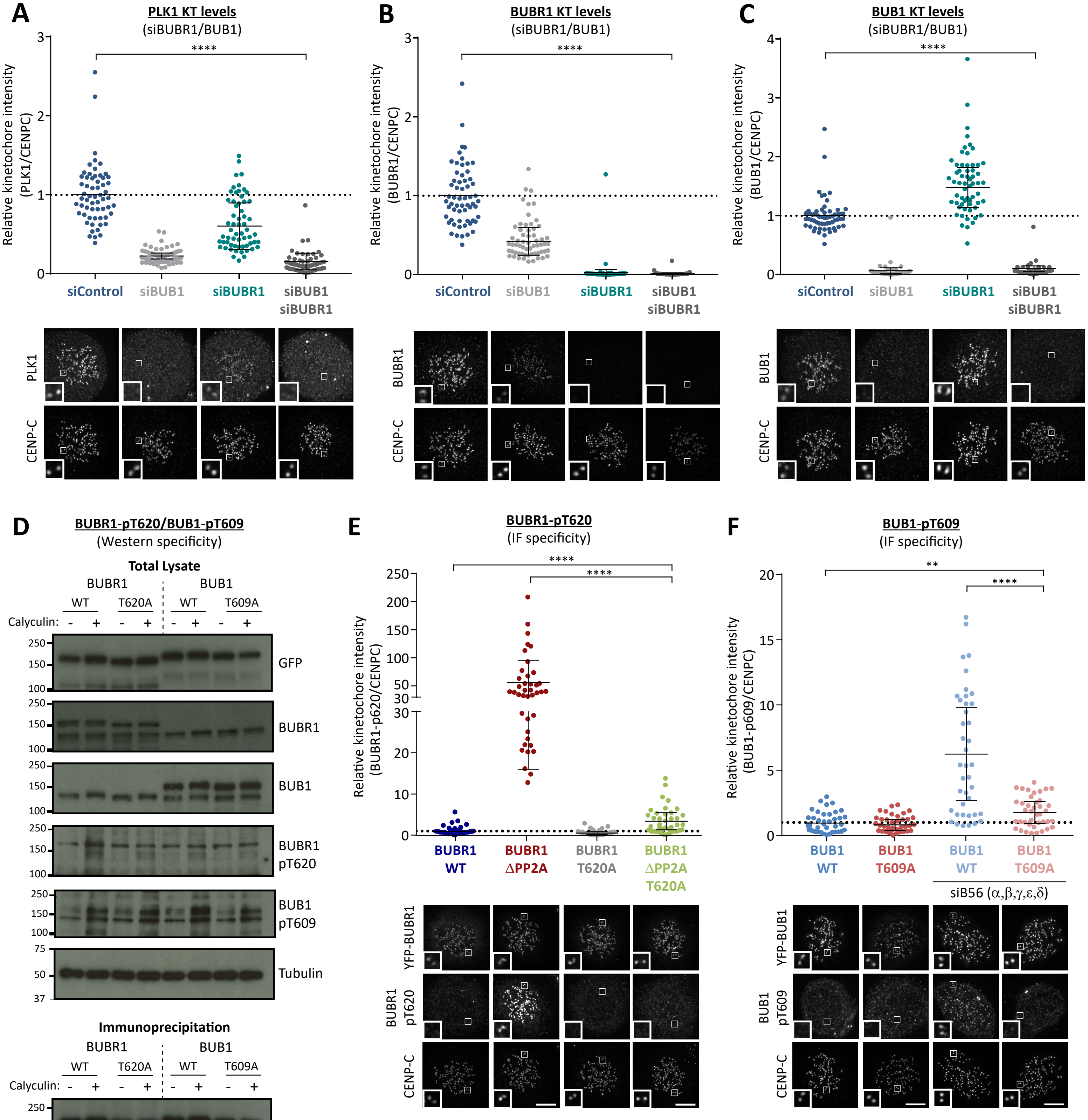

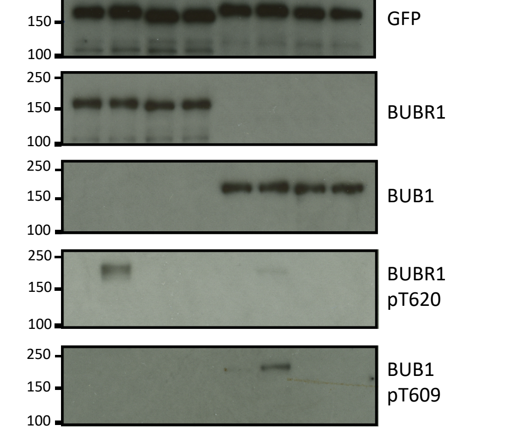

**Figure S1 (related to Figure 1): Testing BUB1/BUBR1 knockdown and phospho-specific antibodies in cells. A-C)** Relative kinetochore levels of PLK1 (A), BUBR1 (B) and BUB1 (C) after siRNA mediated depletion of Bub1, BubR1 or both combined. Graphs show the mean kinetochore intensity of 40-60 cells per condition, 4 experiments. **D)** Mitotic cells expressing exogenous Bub1 and BubR1 constructs were treated with/without 50nM Calyculin for 20 minutes prior to harvesting for lysis to maximally phosphorylate BUB1/BUBR1. YFP-tagged Bub1 and BubR1, mutated in the polo-box binding sites, were immunoprecipitated and blotted with indicated antibodies. Blots from a single experiment were chosen that best represents 3 experiments. **E-F)** Effect of mutating the polo-box binding motifs (BUBR1T620A and BUB1T609A) on antibody detection of BUBR1-pT620 (E) and BUB1-pT609 (F) by immunofluorescence. These experiments were carried out in combination with either mutating the phosphatase binding motif on BUBR1 (E) or siRNA-mediated knock down of all B56 isoforms (F) to elevate BUBR1/BUB1 phosphorylation to maximal levels. The graphs show the mean kinetochore intensity of 40 cells, 4 experiments. For all kinetochore intensity graphs, each dot represents a cell, and the error bars display the variation between the experimental repeats (displayed as ± SD of the experimental means). Example immunofluorescence images were chosen that most closely resemble the mean values in the quantifications. The insets show magnifications of the outlined regions. Scale bars = 5µm. **p<0.01,****p<0.0001

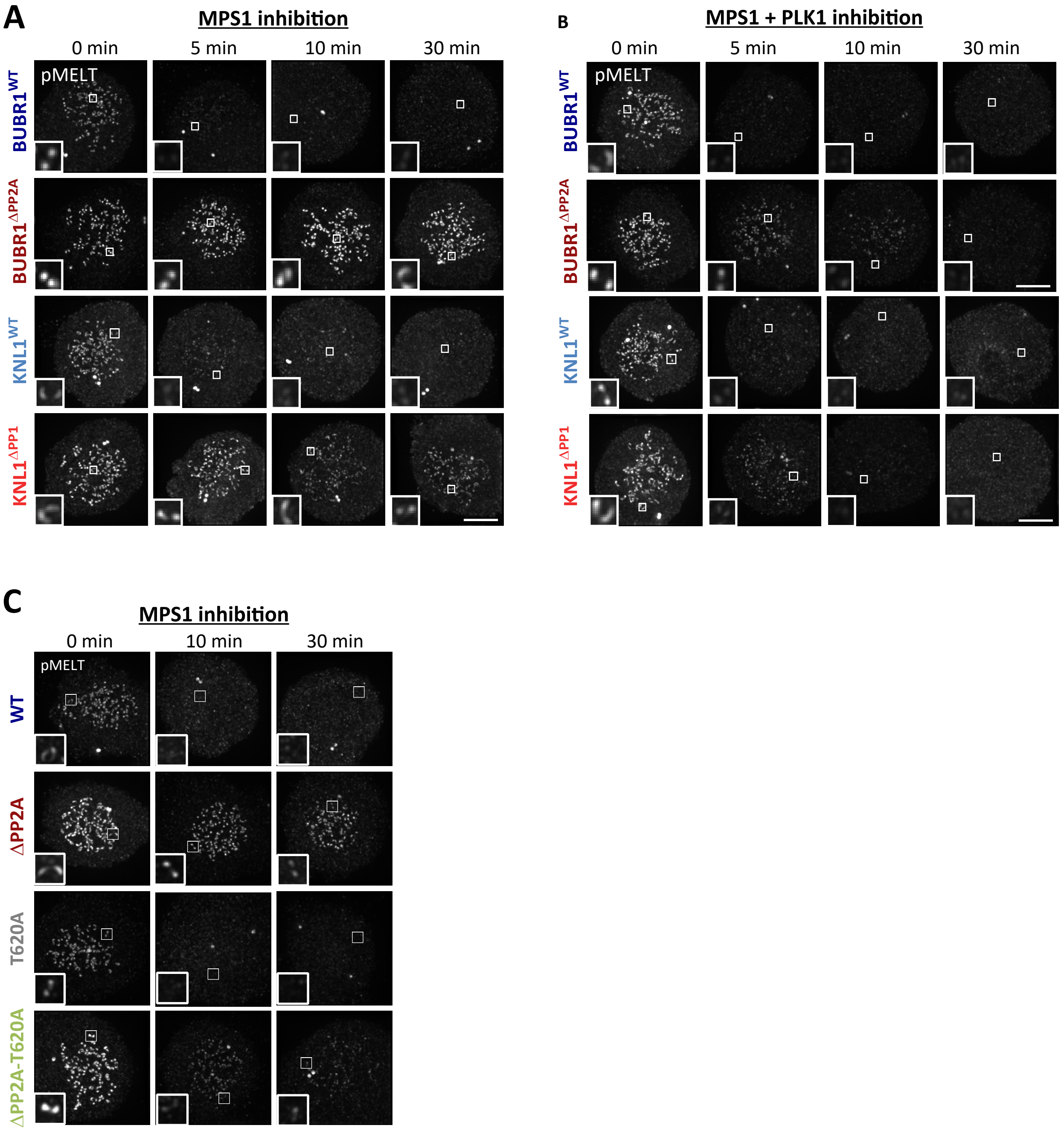


**Figure S2: Example images from kinetochore quantifications in Figure 2. A-C)** Example immunofluorescence images of the kinetochore quantifications shown in Figure 2A (A), 2C (B), 2E (C). The images were chosen that most closely resemble the mean values in the quantifications. The insets show magnifications of the outlined regions. Scale bars = 5µm.

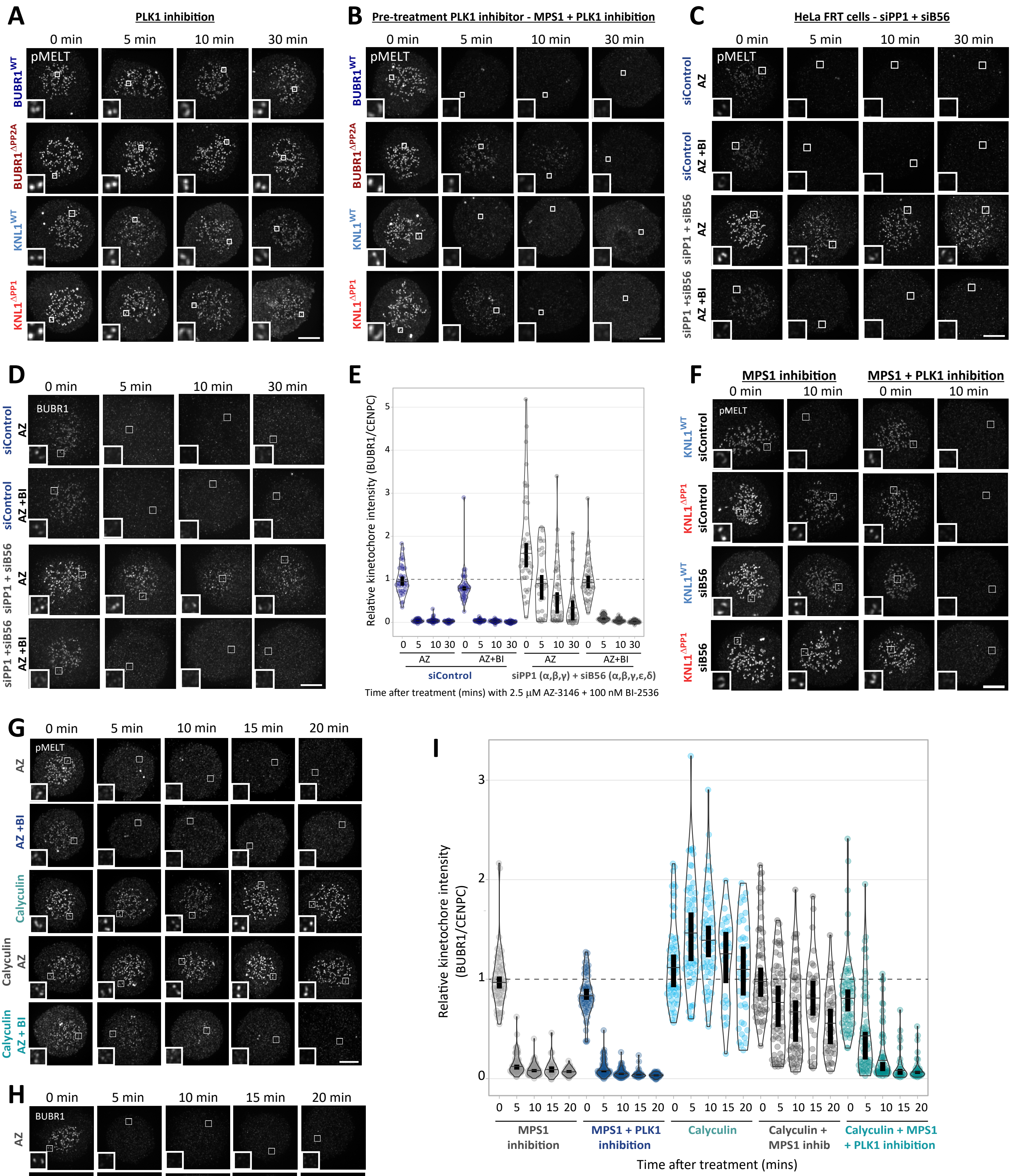

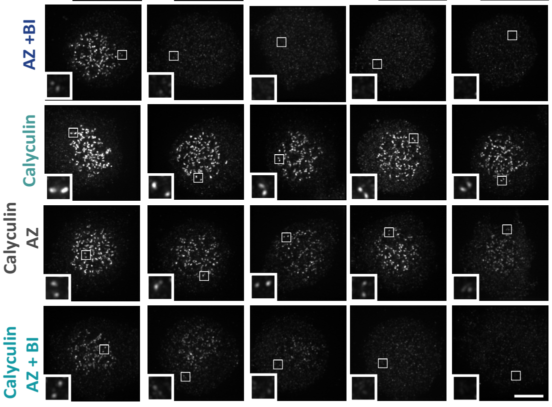

**Figure S3: Example images from kinetochore quantifications in Figure 3. A-C)** Example immunofluorescence images of the kinetochore quantifications shown in Figure 3A (A) 3B (B), 3C (C). **D-E)** Kinetochore BUBR1 images (D) and quantifications (E) from cells treated as in Figure 3C. Graph shows median of 30-40 cells per condition from 3-4 experiments. **F-G)** Example immunofluorescence images of the kinetochore quantifications shown in Figures 3D (F) and 3E (G). **H-I)** Kinetochore BUBR1 images (H) and quantifications (I) from cells treated as in Figure 3E. Graph shows median of 30-60 cells per condition from 6 experiments. MG132 was included in combination with MPS1 inhibitor, in every case, to prevent mitotic exit. For all graphs, each dot represents a cell, and vertical bars show 95% confidence intervals. All images were chosen that most closely resemble the mean values in the quantifications. The insets show magnifications of the outlined regions. Scale bars = 5µm.
